# Multichannel Modulation of Depolarizing and Repolarizing Ion Currents Increases the Positive Rate Dependence of Action Potential Prolongation

**DOI:** 10.1101/2023.01.30.526356

**Authors:** Candido Cabo

## Abstract

A pharmacological approach to prevent reentrant tachycardias is to prolong the action potential duration (APD) of the myocytes that provide the substrate for the arrhythmia. For such an anti-arrhythmic approach to be effective, APD prolongation should be positive rate dependent, that is, it should maximize APD prolongation at the fast excitation rates of tachycardia and minimize APD prolongation at slow excitation rates. APD prolongation by current anti-arrhythmic agents is either reverse (larger APD prolongation at slow rates than at fast rates) or neutral (similar APD prolongation at slow and fast rates), which may not result in an effective anti-arrhythmic action. In this report we show that, in computer models of the human ventricular action potential, the combined modulation of both depolarizing and repolarizing ion currents (I_NaL_, I_CaL_, I_Ks_, I_Kr_ and I_K1_) results in a stronger positive rate dependent APD prolongation than modulation of just repolarizing potassium currents (I_Ks_, I_Kr_ and I_K1_); modulation of a single repolarizing current (I_Kr_) results in reverse rate dependence. A robust positive rate dependent APD prolongation correlates with an acceleration of phase 2 repolarization and a deceleration of phase 3 repolarization, which leads to a triangulation of the action potential shape. A positive rate dependent APD prolongation decreases the repolarization reserve with respect to control at slow excitation rates, which can be managed by interventions that prolong APD at fast excitation rates and shorten APD at slow excitation rates. Feature importance analysis shows that, for both computer models of the action potential, I_CaL_ and I_K1_ are the most important ion currents to achieve a positive rate dependent APD prolongation. In conclusion, multichannel modulation of depolarizing and repolarizing ion currents, with ion channel activators and blockers, results in a robust APD prolongation at fast excitation rates, which should be anti-arrhythmic, while minimizing APD prolongation at slow heart rates, which should reduce potential pro-arrhythmic risks.

## INTRODUCTION

A strategy for the treatment of reentrant tachyarrhythmias is the prolongation of the action potential duration (APD) using potassium channel blockers (Peters et al 2000; Tamargo et al 2004). However, clinical trials have shown that agents that selectively block potassium ion channels (like I_Kr_, the rapid delayed rectifier potassium channel) are not effective at preventing initiation and maintenance of reentrant arrhythmias in post-myocardial infarction patients (Waldo et al. 1996; Bloch-Thomsen 1998; Køber et al. 2000). APD prolongation with specific I_Kr_ blockers is reverse rate dependent, that is, APD prolongation is greater at slow excitation rates than at the fast excitation rates of ventricular tachycardia. That is far from optimal, not only because APD may not be sufficiently prolonged at the fast rates typical of ventricular tachycardia (Hondeghem and Snyders 1990), but also because APD prolongation at slow rates may result in drug-induced LQT syndrome, trigger early afterdepolarizations and have a pro-arrhythmic effect (Kannankeril et al. 2010). For APD prolongation to be anti-arrhythmic, APD prolongation should exhibit a positive rate dependent response, that is, it should maximize APD prolongation at fast excitation rates and minimize APD prolongation at slow excitation rates.

Consistent with experimental and clinical evidence, we showed in an earlier report (Cabo 2022) that, in computer models of the human ventricular action potential, APD prolongation by blocking selectively a specific potassium ion current type results in reverse or a marginal positive rate dependence (Cabo 2022). In contrast, APD prolongation by the combined modulation of several potassium ion currents (I_Ks_, I_Kr_ and I_K1_) can result in a robust positive rate dependence, suggesting the potential of multichannel pharmacology as an antiarrhythmic strategy (Cabo 2022). There is clinical evidence that multichannel pharmacology results in more effective antiarrhythmic agents than pharmacology that selectively modulates a specific ion current. For example, amiodarone, an agent that is effective against a variety of arrhythmias, prolongs the APD by blocking sodium and potassium currents and has an attenuated reverse rate dependence response when compared to agents that block selectively a specific ion current (Hondeghem and Snyders 1990; Dorian and Newman 2000). In this report we further investigate how multichannel pharmacology can result in a robust positive rate dependent APD prolongation. We hypothesized that the combined modulation of depolarizing (I_NaL_ and I_CaL_) and repolarizing currents (I_Ks_, I_Kr_ and I_K1_) can further increase the positive rate dependence of APD prolongation. As before, we used computational models of the human ventricular action potential in combination with optimization algorithms to investigate which interventions may result in prolongation of APD with a positive rate dependence, and to further understand how the shape of the action potential relates to its rate dependence. We also investigated the relative importance of ion currents to achieve a positive rate dependent APD prolongation.

## METHODS

### Computer models of the action potential

We simulated the cardiac action potential using the ORd (O’Hara et al. 2011) and the ToR-ORd (Tomek et al. 2019) models of a human ventricular epicardial cell. Both models are publicly available: the ORd model was downloaded from the Rudy Lab web site (https://rudylab.wustl.edu/code-downloads/) and the ToR-ORd model was downloaded from the CellML repository (www.cellml.org). The ToR-ORd model builds on the structure of the ORd model, but the formulation of several currents that determine the rate dependence of the action potential, like I_CaL_, I_Kr_ and I_K1_, is different (Tomek et al. 2019).

We investigated the rate dependence of the action potential models by modulating the maximum conductance of two depolarizing currents: the late sodium current (I_NaL_; range: 0-2x control) and the L-type calcium current (I_CaL_; range: 0.5-1.5x control); and three repolarizing currents: the slow delayed rectifier potassium current (I_Ks_; range: 0-20x control), the rapid delayed rectifier potassium current (I_Kr_; range: 0.5-2x control) and the inward rectifier potassium current (I_K1_; range: 0.2-2x control). Action potentials were initiated with a depolarizing current with a strength 1.5x the stimulation threshold. We report measurements on action potentials that were calculated after 30 minutes of stimulation to achieve steady-state.

### Action potential features

To quantify and compare different action potential shapes we calculated the average ion current during phase 2 (I_ion, phase2_) and during phase 3 (I_ion, phase3_) as well as their ratio (I_ion, phase3_ / I_ion, phase2_). The phases of the action potential were quantified as described in an earlier report (Cabo 2022). In short, phase 1 begins at the time of action potential depolarization and it ends at the time repolarization starts, which is when the total ion current becomes positive (Cabo 2022, Figure 1). Phase 2 starts when phase 1 ends, and it ends when I_K1_ raises to 10% of its peak (Cabo 2022). In the ORd model the end of phase 2 occurs when the membrane repolarizes to −39 mV. In the ToR-ORd model the end of phase 2 occurs when the membrane repolarizes to −34 mV. Phase 3 starts at the end of phase 2, and it ends when the action potential repolarizes by 90% of the action potential amplitude from its maximum depolarization potential.

**Figure 1.**
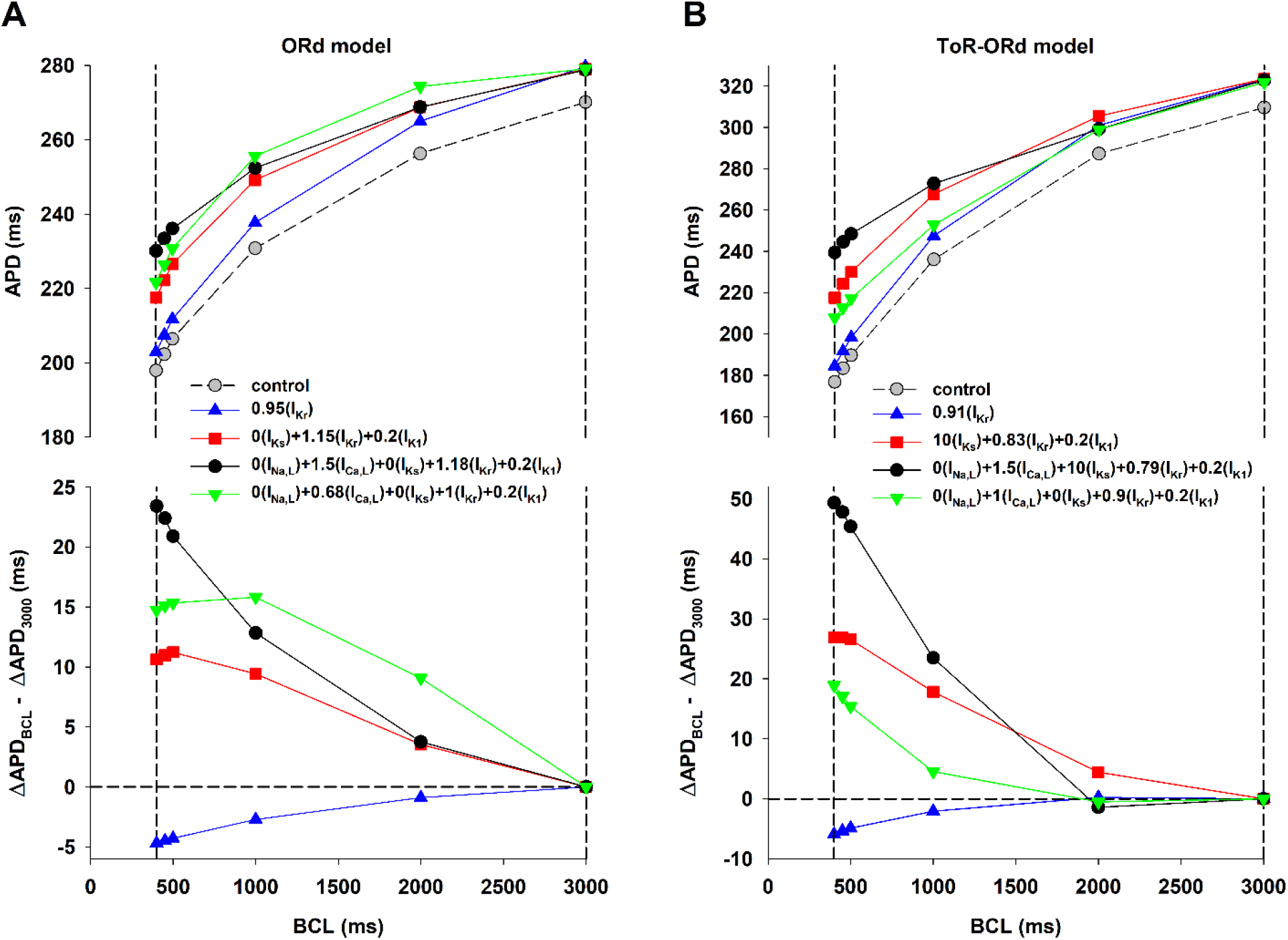
Rate dependence of APD prolongation with the ORd (panel A) and ToR-ORd (panel B) models. Panel A (top): APD rate dependence during control (gray circles), and for four interventions that prolonged the control APD to 280 ms when BCL = 3000 ms, using the ORd model. The vertical dashed lines indicate BCL = 400 and 3000 ms respectively. Panel A (bottom): Difference between APD prolongation (with respect to control) for each BCL and APD prolongation when BCL = 3000 ms for the four interventions in Panel A (top). Positive values indicate positive rate dependence and negative values indicate reverse rate dependence. Panel B shows similar results for four interventions that prolong the control APD to 320 ms, when BCL = 3000 ms, using the ToR-ORd model. The format is the same as in panel A. See text for detailed description.

### Estimation of the repolarization reserve

We estimated the repolarization reserve of a baseline action potential by quantifying the prolongation of the APD upon application of a constant depolarizing current of −0.1pA/pF during the action potential (Varro and Baczko 2011). This can be done experimentally for example by increasing the late sodium current (I_NaL_) with veratrine and anemonia sulcata toxin (ATX II) (Varro and Baczko 2011). With that protocol, a larger prolongation of APD with respect to the baseline APD implies a smaller repolarization reserve and a higher risk of triggered arrhythmias.

### Particle Swarm Optimization algorithm

When using specific potassium channel blockers to prolong the action potential, the prolongation of the action potential (with respect to control) at long cycle lengths (ΔAPD_long_) is generally larger than the prolongation of the action potential at short cycle lengths (ΔAPD_short_), resulting in reverse rate dependence (ΔAPD_long_ > ΔAPD_short_). As before (Cabo 2022), we used the Particle Swarm Optimization (PSO) algorithm (Kennedy and Eberhart, 1995) to find the optimal combination of maximum conductance of I_NaL_, I_CaL_, I_Ks_, I_Kr_ and I_K1_ to minimize the difference between ΔAPD_long_ and ΔAPD_short_ while achieving a given APD prolongation. We used an implementation of the PSO algorithm publicly available in the Github repository (https://github.com/kkentzo/pso). Minimizing ΔAPD_long_-ΔAPD_short_ should result in an attenuation of reverse rate dependence or achieving positive rate dependence if ΔAPD_long_ < ΔAPD_short_. In the simulations presented here, for both the ORd and ToR-ORd model, the long cycle length was BCL = 3000 ms, and the short cycle length was BCL = 400 ms. The input of the PSO optimization algorithm was the APD goal and the allowed range of variation (minimum and maximum values) of ion channel conductances.

### Backward Feature Elimination

We used a backward feature elimination procedure to investigate the relative contribution of each ion current to the action potential positive rate dependence response. After applying the PSO optimization to find the combination of maximum conductance of I_NaL_, I_CaL_, I_Ks_, I_Kr_ and I_K1_ to maximize a positive rate dependence response, optimization was applied to the five possible subsets of four currents (i.e., [I_CaL_, I_Ks_, I_Kr_ and I_K1_], [I_NaL_, I_Ks_, I_Kr_ and I_K1_], [I_NaL_, I_CaL_, I_Kr_ and I_K1_], [I_NaL_, I_CaL_, I_Ks_ and I_K1_], [I_NaL_, I_CaL_, I_Ks_ and I_Kr_]). The subset resulting in the larger positive rate dependence response after PSO optimization was selected for the next step in the elimination procedure. The ion current not present in the selected subset was the current that contributed less to the positive rate dependence response and it was consequently eliminated. This process of elimination was repeated until only one ion current was left.

## RESULTS

### Positive rate dependence with multi-channel pharmacology

Figure 1A (top) shows the APD rate dependence during control (gray circles, dashed line), and for four interventions that prolonged the control APD to 280 ms at BCL = 3000 ms, using the ORd model. While APD prolongation at BCL = 3000 ms (right vertical dashed line) is the same for all interventions (~10 ms with respect to control), APD prolongation for shorter BCLs was markedly different (left vertical dashed line), indicating a different APD rate dependence. Figure 1A (bottom) shows how APD prolongation with respect to control at different BCLs compares to APD prolongation at BCL = 3000 ms for the four interventions. A positive value indicates that APD prolongation at a given BCL is larger that APD prolongation at 3000 ms, and shows a positive rate dependence; a negative value indicates a reverse rate dependence. The optimal combinations of ion channel activators and/or blockers were obtained using the PSO optimization algorithm described in the Methods.

It is well established experimentally and computationally that prolonging APD by blocking I_Kr_ (Figure 1A, blue triangles up) results in a reverse rate dependent response (Dorian and Newman 2000; Virag et al. 2009; Banyasz et al. 2009; Barandi et al 2010; Cabo 2022). However, by modulating repolarizing potassium currents (I_Ks_, I_Kr_, and I_K1_), it is possible to achieve a positive rate dependent response (Figure 1, red squares) (Cabo 2022). Moreover, Figure 1A shows that modulation of both repolarizing currents (I_Ks_, I_Kr_, and I_K1_) and depolarizing currents (I_NaL_ and I_CaL_) (Figure 1, black circles) drastically enhances the positive rate dependent APD prolongation at faster rates (i.e., BCLs < 500 ms). Figure 1A also shows that modulation of depolarizing and repolarizing currents with ion channel activators and blockers (Figure 1, black circles) results in a more robust positive rate dependent APD prolongation than modulation of the same currents with ion channel blockers at BCLs < 500 ms (Figure 1, green triangles down).

Figure 1B shows similar results for interventions that prolong APD to 320 ms using the ToR-ORd model (a prolongation of ~10 ms with respect to control). Note that there are quantitative differences between both models, a consequence of the different formulation of I_CaL_, I_Kr_ and I_K1_. For example, APD during control at BCL = 3000 ms is larger for the ToR-ORd model (310 ms) than for the ORd model (270 ms). In contrast, the control APD of the ToR-ORd model at BCL = 400 ms is shorter than that of the ORd model, which indicates a larger APD rate adaptation of the ToR-ORd model. The larger y-axis values in Figure 1A and 1B (bottom) also show that the positive rate dependence response is more robust for the ToR-ORd model than for the ORd model.

Despite quantitative differences between the models, the results in Figure 1 show that modulating both depolarizing and repolarizing ion currents results in a stronger positive rate dependence than modulation of just repolarizing potassium currents. Moreover, modulation of ion currents with ion channel activators and blockers produces a more robust positive rate dependent APD prolongation than modulation of the same ion currents with only ion channel blockers. Modulation of a single repolarizing current (I_Kr_) results in reverse rate dependence.

### Action potential shape and its effect on a positive rate dependence

Figure 2A (top) shows action potentials for three of the interventions in Figure 1A that resulted in APD = 280 ms with BCL = 3000 ms, a prolongation of about 10 ms the control APD, using the ORd model. The action potentials are superimposed to make it easier to compare their shapes. Figure 2A (bottom) shows the corresponding total ion currents (I_ion_) during the action potentials. Figure 2A also shows the duration of phase 2 and phase 3 for the different action potentials (red and blue vertical dashed lines). Phase 2 starts at the leftmost vertical dashed line (Figure 2A) and ends when action potentials repolarize to −39 mV (red and blue vertical dashed lines). The end of phase 2 for the action potentials with multichannel modulation (black and red action potentials in Figure 2A) is the same. Phase 3 begins at the end of phase 2 and ends at the time of 90% repolarization (rightmost vertical dashed line in Figure 2A); the end of phase 3 is the same for all action potentials.

**Figure 2.**
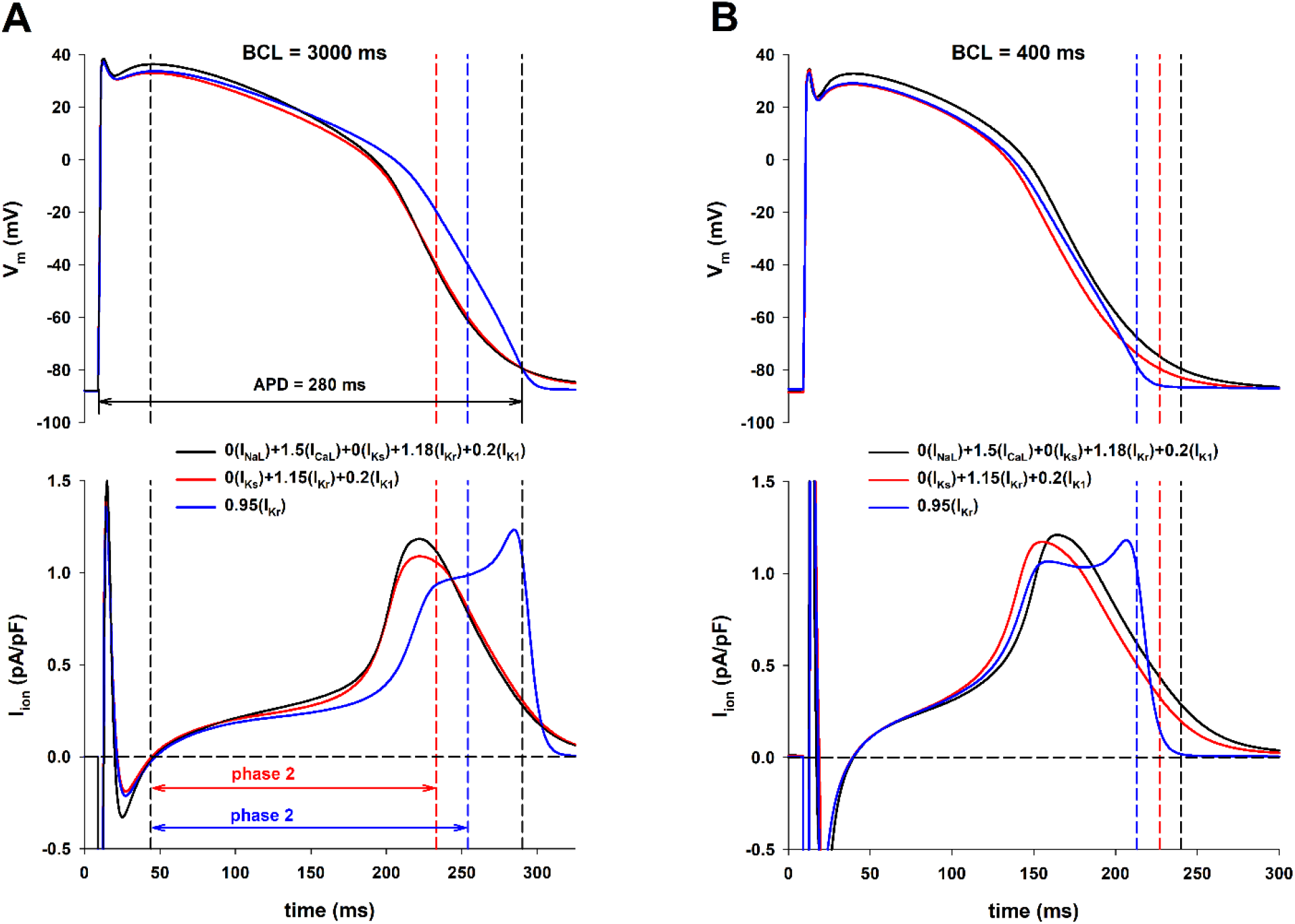
Panel A: Action potentials for different interventions that prolong the action potential to 280 ms when BCL = 3000 ms (top) with the corresponding total ion currents (bottom), using the ORd model. The leftmost vertical black dashed line indicates the beginning of phase 2 of the action potential. The vertical red dashed line indicates the end of phase 2 for the (red and black) action potentials with a positive rate dependence. The vertical blue dashed line indicates the end of phase 2 for the (blue) action potential with a reverse rate dependence. The rightmost vertical black dashed line indicates the time of 90% repolarization, which is the same for the three action potentials. Panel B: Action potentials when BCL = 400 ms for the interventions in panel A (top) with the corresponding total ion currents (bottom). The dashed (color coded) vertical lines indicate the time of 90% repolarization for the three interventions. See text for detailed description.

Interventions that result in a positive rate dependence (red and black lines in Figure 2A, bottom) have a larger I_ion_ during early action potential repolarization than the intervention that results in reverse rate dependence (blue line in Figure 2A, bottom). In contrast, during late repolarization of the action potential, I_ion_ is larger for the intervention that results in reverse rate dependence than for interventions that result in a positive rate dependence. This is reflected in a shorter phase 2 (and consequently a longer phase 3) for interventions that result in positive rate dependence (Figure 2A, bottom).

The duration of phase 2 and phase 3 for the two interventions that result in a positive rate dependence is about the same (red and black action potentials in Figure 2A). But note that I_ion_ during phase 2 is larger for the intervention that modulated *both* depolarizing and repolarizing currents and that resulted in a larger positive rate dependent response (compare black and red action potentials in Figure 2A, bottom, during phase 2); this is a consequence of the larger depolarization produced by the 50% increase in I_CaL_ at the beginning of phase 2. The total I_ion_ for the interventions that result in a positive rate dependence (black and red action potentials in Figure 2A, bottom) during phase 3 is about the same, which is consistent with the fact that I_K1_, the largest current during late repolarization, is blocked by the same amount (80%) for both interventions.

Figure 2B (top) shows the corresponding action potentials for the three interventions when BCL = 400 ms. The vertical dashed lines indicate the time of 90% repolarization for the different action potentials to illustrate the differences in APD for the three interventions at the faster stimulation rate. Figure 2B (bottom) shows the corresponding total ion currents. The results are qualitatively similar to those in Figure 2A, that is, positive rate dependence is associated with faster repolarization during phase 2 and slower repolarization during phase 3 of the action potential. Overall, the results in Figure 2 show that, in the ORd model, a positive rate dependent response is associated with a larger I_ion_ (i.e., a faster repolarization) during phase 2 and a smaller I_ion_ (i.e., a slower repolarization) during phase 3 of the action potential.

Figure 3 (top) shows action potentials for three of the interventions in Figure 1B with BCL = 3000 ms (Figure 3A) and BCL = 400 ms (Figure 3B), using the ToR-ORd model. Figure 3 (bottom) shows the corresponding total ion currents (I_ion_) during the action potentials. The figure format is the same as in Figure 2. Note that in spite of the quantitative differences in the formulation of the models, the results obtained with the ORd model (Figure 2) and the ToR-ORd model (Figure 3) are similar: a positive rate dependent response is associated with a larger I_ion_ (i.e., a faster repolarization) during phase 2 and a smaller I_ion_ (i.e., a slower repolarization) during phase 3 of the action potential.

**Figure 3.**
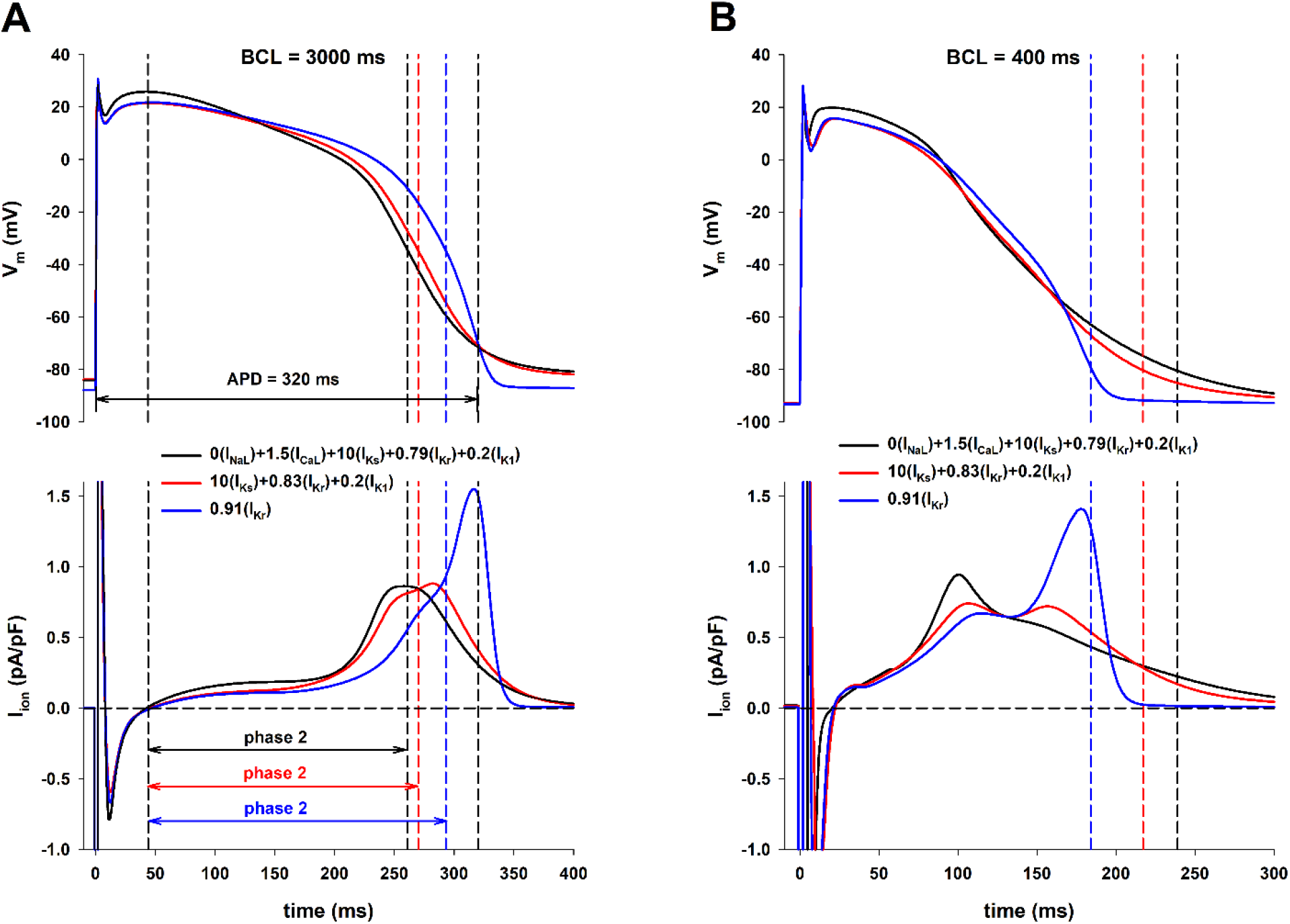
Panel A: Action potentials for different interventions that prolong the action potential to 320 ms when BCL = 3000 ms (top) with the corresponding total ion currents (bottom). The leftmost vertical black dashed line indicates the beginning of phase 2 of the action potential. The vertical red/black dashed line indicate the end of phase 2 for the red/black action potential with a positive rate dependence. The vertical blue dashed line indicates the end of phase 2 for the blue action potential with a reverse rate dependence. The rightmost vertical black dashed line indicates the time of 90% repolarization, which is the same for the three action potentials. Panel B: Action potentials when BCL = 400 ms for the interventions in panel A (top) with the corresponding total ion currents (bottom). The dashed (color coded) vertical lines indicate the time of 90% repolarization for the three interventions. See text for detailed description.

Figure 4 shows a summary of average I_ion_ during phase 2 and phase 3 as well as their ratio, for different BCLs, for the ORd model (Figure 4A) and the ToR-ORd model (Figure 4B). APDs for all interventions are the same only with BCL = 3000 ms; for shorter BCLs, APDs varied because different interventions have a different rate dependence (Figure 1). For all BCLs, interventions with a strong positive rate dependence (black circles and red squares) exhibit a larger average I_ion_ during phase 2 (top panel), a smaller average I_ion_ during phase 3 (middle panel), and a smaller I_ion, phase3_ / I_ion, phase2_ ratio (bottom panel) than for the intervention with reverse rate dependence (blue triangles up). Figure 4 also shows that a faster stimulation rate (i.e., a shorter BCL) results in a robust increase of average I_ion_ during phase 2, with a moderate (in the ToR-ORd model) or marginal decrease (in the ORd model) of average I_ion_ during phase 3. Consequently, most of the shortening of the action potential with rate in both models occurs as a result of a shortening of phase 2, with phase 3 remaining relatively constant or slightly increased.

**Figure 4.**
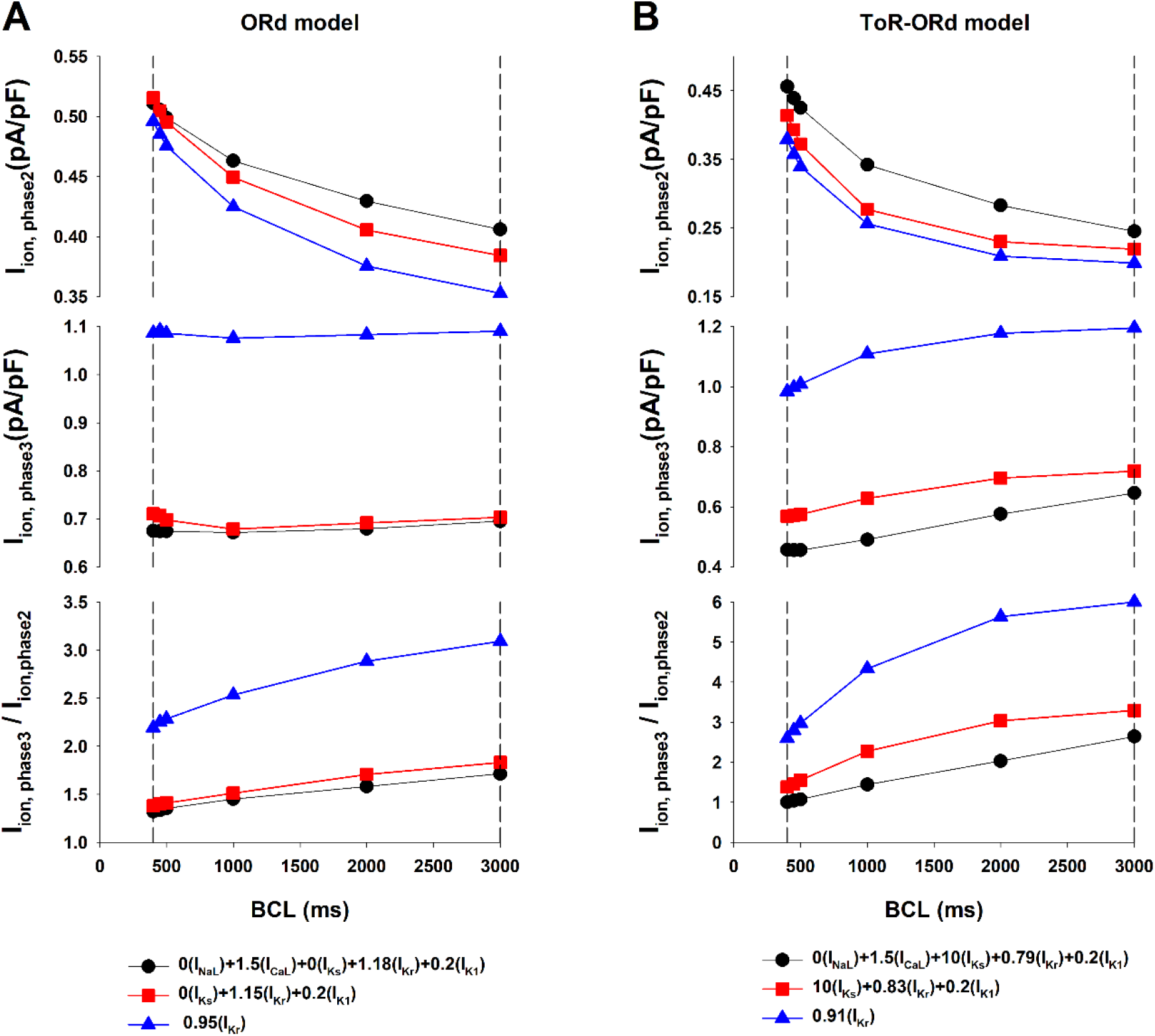
Average I_ion_ during phase 2 (top) and phase 3 (middle) of the action potential as well as their ratio I_ion, phase3_ / I_ion, phase2_ (bottom), for different BCLs, using the ORd model (panel A) and the ToR-ORd model (panel B). See text for detailed description.

Interventions that prolong the action potential with a positive rate dependence result in a smaller I_ion, phase3_ / I_ion, phase2_ ratio. Average I_ion_ over a time interval is the average slope of action potential repolarization during that interval (Cabo 2022); therefore, a smaller I_ion, phase3_ / I_ion, phase2_ ratio indicates a triangulation of the action potential shape (Figures 2 and 3). Action potential triangulation may result in the development of triggered arrhythmias (Hondeghem et al 2001; Shah and Hondeghem 2005; Kannankeril et al 2010). To compare the pro-arrhythmic potential of the interventions in Figures 1–4 we estimated their repolarization reserve with BCL = 3000 ms (Figure 5). The figure shows that interventions that result in a positive rate dependence (Figure 5A and 5B, bars #2 and #3) have a decreased repolarization reserve with respect to control (Figure 5A and 5B, bar #1). For the ORd model, there is no difference in the repolarization reserve for the intervention that modulates both depolarizing and repolarizing currents (Figure 5A, bar #2) and the intervention that modulates only repolarizing currents (Figure 5A, bar #3). In contrast, for the ToR-ORd model, the intervention with a stronger positive rate dependent response (Figure 5B, bar #2) has a smaller repolarization reserve that the intervention with a smaller positive rate dependent response (Figure 5B, bar #3).

**Figure 5.**
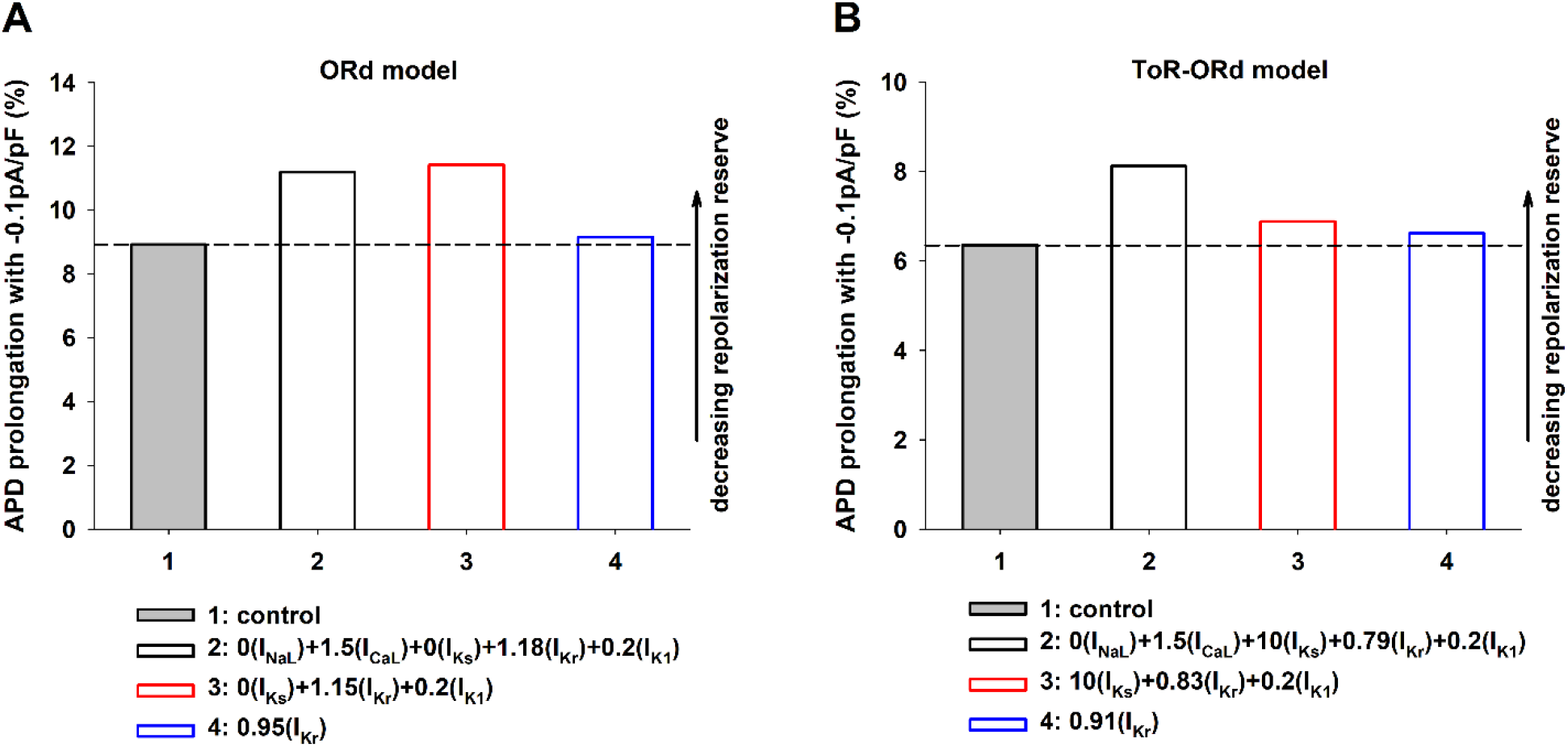
Percentage prolongation of APD when a constant depolarizing current of −0.1pA/pF is applied during the action potential when BCL = 3000 ms for the control action potential (gray bar, dashed horizontal line) and for the three interventions in Figures 2–4. Panel A: Interventions that result in a positive rate dependence (bars #2 and #3) and a reverse rate dependence (bar#4) using the ORd model. Panel B: Interventions that result in positive rate dependence (bars #2 and #3) and reverse rate dependence (bar#4) using the ToR-ORd model. For interventions with positive rate dependence (bar#2 and bar#3 in Panels A and B), APD prolongation is above the dashed line indicating that the repolarization reserve is decreased with respect to control. See text for detailed description.

### Relative importance of ion currents to achieve a positive rate dependent APD prolongation

The previous results show that the combined modulation of depolarizing currents (I_NaL_ and I_CaL_) and repolarizing currents (I_Ks_, I_Kr_, and I_K1_) results in a robust positive rate dependence (Figure 1A and 1B, black circles). We used the backward feature elimination procedure to investigate the relative importance of each current to the positive rate dependent APD prolongation to 280 ms with BCL = 3000 ms, using the ORd model (Figure 1A, black circles). Figure 6 shows the four steps in the procedure for the ORd model: at each step the current that contributes less to the positive rate dependence is eliminated. Step 1 shows ΔAPD_BCL=400ms_ - ΔAPD_BCL=3000ms_ to estimate the action potential rate dependence response when all depolarizing and repolarizing ion currents are modulated. From step 1 to step 2, I_Ks_ is eliminated because that is the ion current that contributes less to the positive rate dependence response; however, the elimination of I_Ks_ results in a decreased positive rate dependence response. I_NaL_ and I_Kr_ are eliminated in steps 3 and 4, with the consequent decrease in positive rate dependence. The backward feature elimination procedure identifies I_CaL_ and I_K1_ as the ion currents that are more important for a positive rate dependence. Figure 6 also shows that the modulation of each channel individually does not result in an action potential with a positive rate dependence.

**Figure 6.**
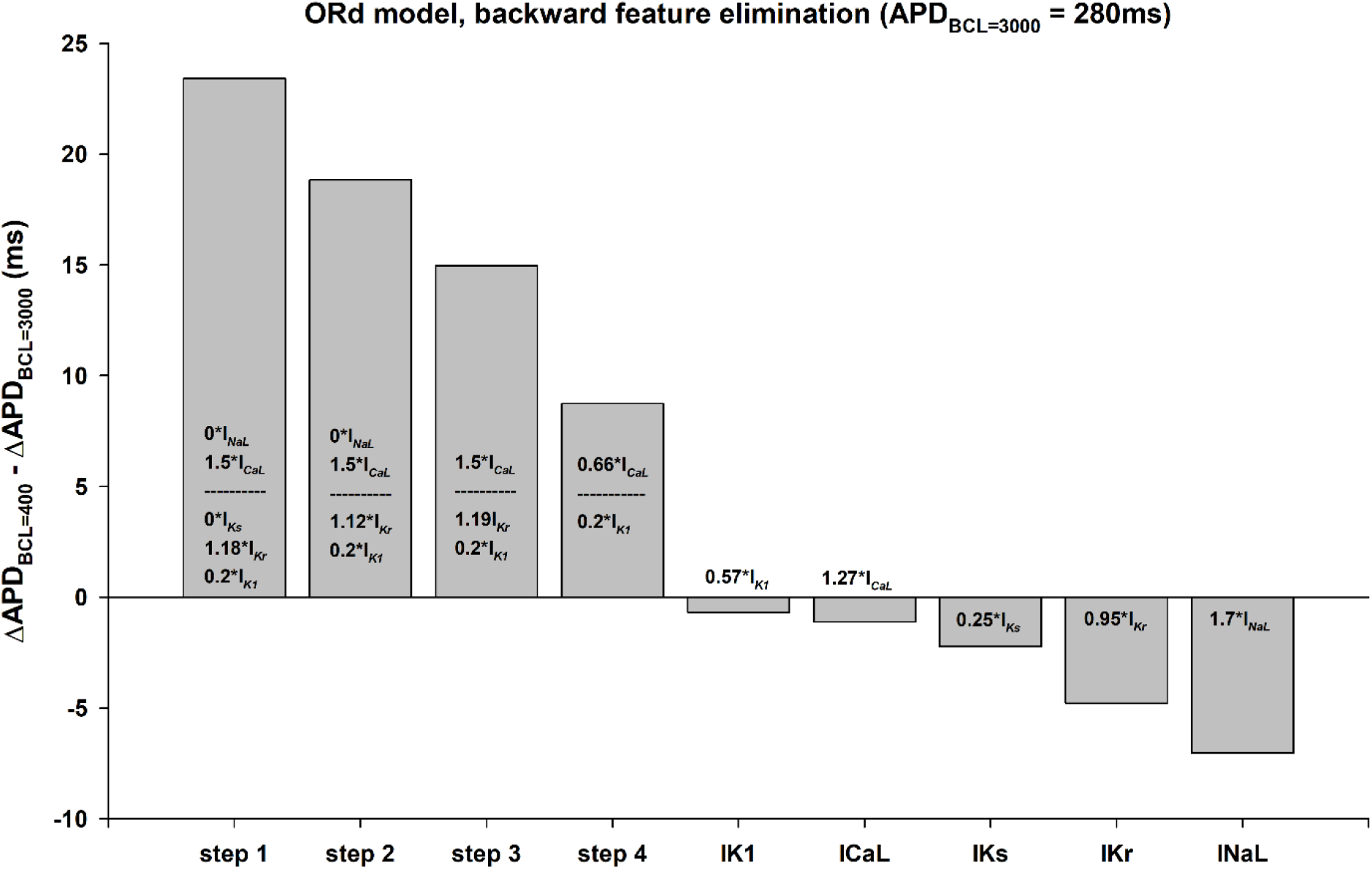
Relative importance of each ion current to a positive rate dependent APD prolongation to 280 ms, when BCL = 3000 ms, using the ORd model. The figure shows the four steps in the backward feature elimination procedure for the intervention that modulates both depolarizing and repolarizing ion currents (0I_NaL_+1.5I_CaL_+0I_Ks_+1.18I_Kr_+0.2I_K1_) in Figure 1A (black circles). At each step, the ion current that contributes less to the positive rate dependence is eliminated. The interventions shown inside the bars are the ion current modulations that maximize the positive rate dependence after a current is eliminated. The y-axis shows the difference between APD prolongation (with respect to control) when BCL = 400 ms and APD prolongation when BCL = 3000 ms: a positive value indicates positive rate dependence. The figure also shows that the modulation of each channel individually does not result in a positive rate dependence APD prolongation to 280 ms in the ORd model. See text for detailed description.

Figure 7 shows the results of using the backward feature elimination procedure to investigate the relative importance of each ion current to the positive rate dependent APD prolongation to 320 ms with BCL = 3000 ms, using the ToR-ORd model (Figure 1B, black circles). The two most important currents for a positive rate dependence response, when the backward feature elimination procedure reaches step 4, are I_CaL_ and I_K1_. As we mentioned earlier, the ToR-ORd model shows a stronger positive rate dependence than the ORd model. Indeed, individual modulation of I_CaL_ and I_K1_ also results in a, somehow diminished, positive rate dependence response, which did not occur for the ORd model. Figure 7 does not show the effect of I_Ks_ block on positive rate dependence because, given the small contribution of I_Ks_ to action potential repolarization in the ToR-ORd model, complete block of I_Ks_ was not enough to prolong APD to 320 ms.

**Figure 7.**
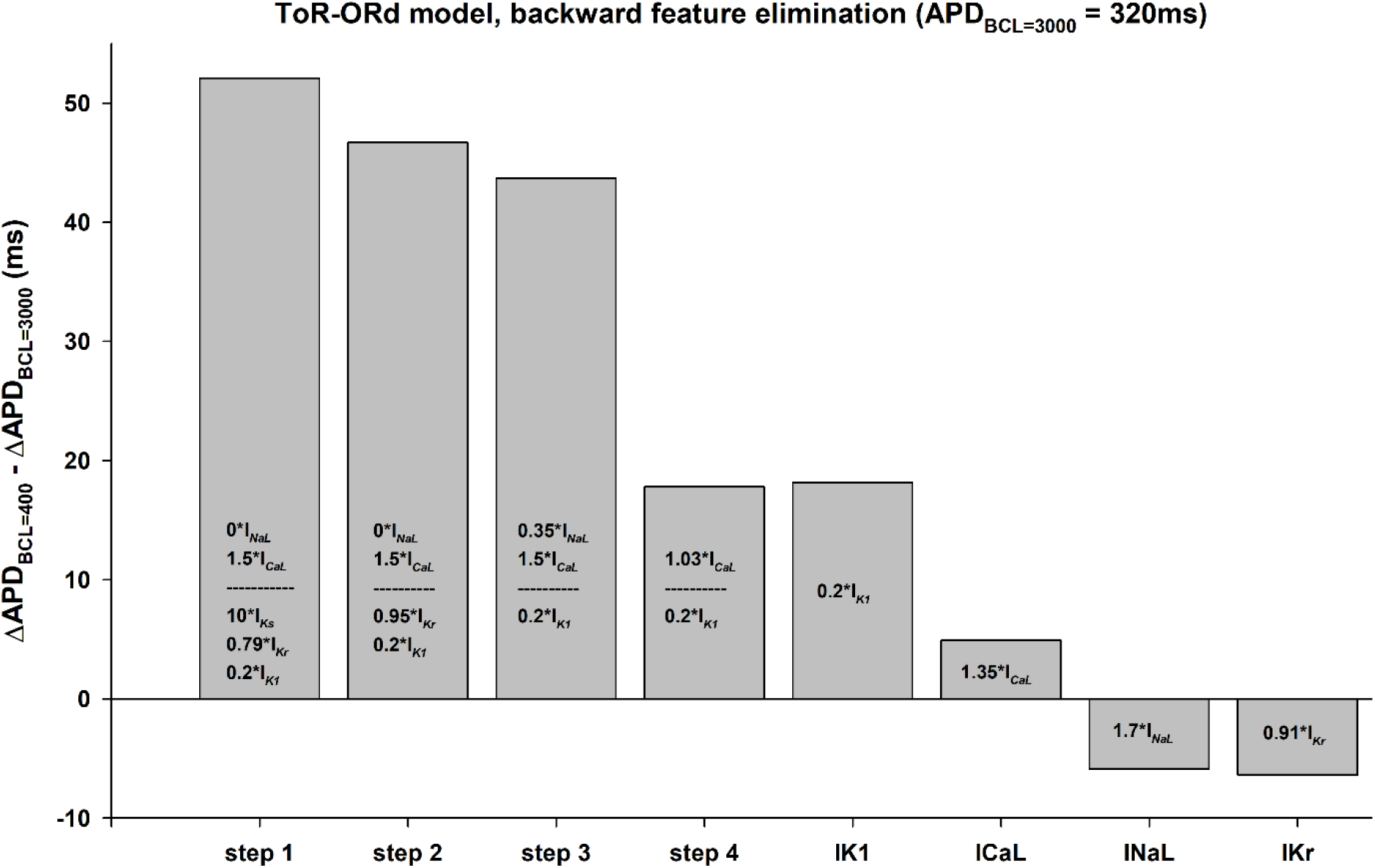
Relative importance of each ion current to a positive rate dependent APD prolongation to 320 ms, when BCL = 3000 ms, using the ToR-ORd model. The figure shows the four steps in the backward feature elimination procedure for the intervention that modulates both depolarizing and repolarizing ion currents (0I_NaL_+1.5I_CaL_+10I_Ks_+0.79I_Kr_+0.2I_K1_) in Figure 1B (black circles). At each step, the ion current that contributes less to the positive rate dependence is eliminated. The interventions shown inside the bars are the ion current modulations that maximize the positive rate dependence after a current is eliminated. The y-axis shows the difference between APD prolongation (with respect to control) when BCL = 400 ms and APD prolongation when BCL = 3000 ms: a positive value indicates positive rate dependence. The figure also shows that specific modulation of I_CaL_ or I_K1_ also results in positive rate dependence in the ToR-ORd model. See text for detailed description.

### Action potential duration prolongation of different magnitudes with a positive rate dependence

The modulation of depolarizing and repolarizing ion currents in Figures 1–7 produced an APD prolongation of about 10 ms with respect to control at a BCL = 3000 ms. Varying the modulation of I_Kr_ while keeping modulation of the other ion currents at the optimal values that resulted in a positive rate dependence makes it possible to produce different amounts of APD prolongation (Figure 8). Figure 8A (top) shows the rate dependent responses of control (dashed black line) and 4 interventions that result in positive rate dependence for different modulations (activations) of I_Kr_ resulting in different APD prolongations in the ORd model. Indeed, it is possible to prolong APD with BCL = 400 ms with no APD prolongation (X=1.23I_Kr_, blue triangles up in Figure 8A, top) or even with a APD shortening with BCL = 3000 ms (X=1.3I_Kr_, red triangles down in Figure 8A, top). Figure 8A (bottom) shows that larger APD prolongations results in a decrease in the positive rate dependent response. For example, the larger APD prolongation (X=1.13I_Kr_, ochre squares in Figure 8A, top) leads to the smaller rate dependent response (X=1.13I_Kr_, ochre squares in Figure 8A, bottom). Figure 8B shows similar results for the ToR-ORd model.

**Figure 8.**
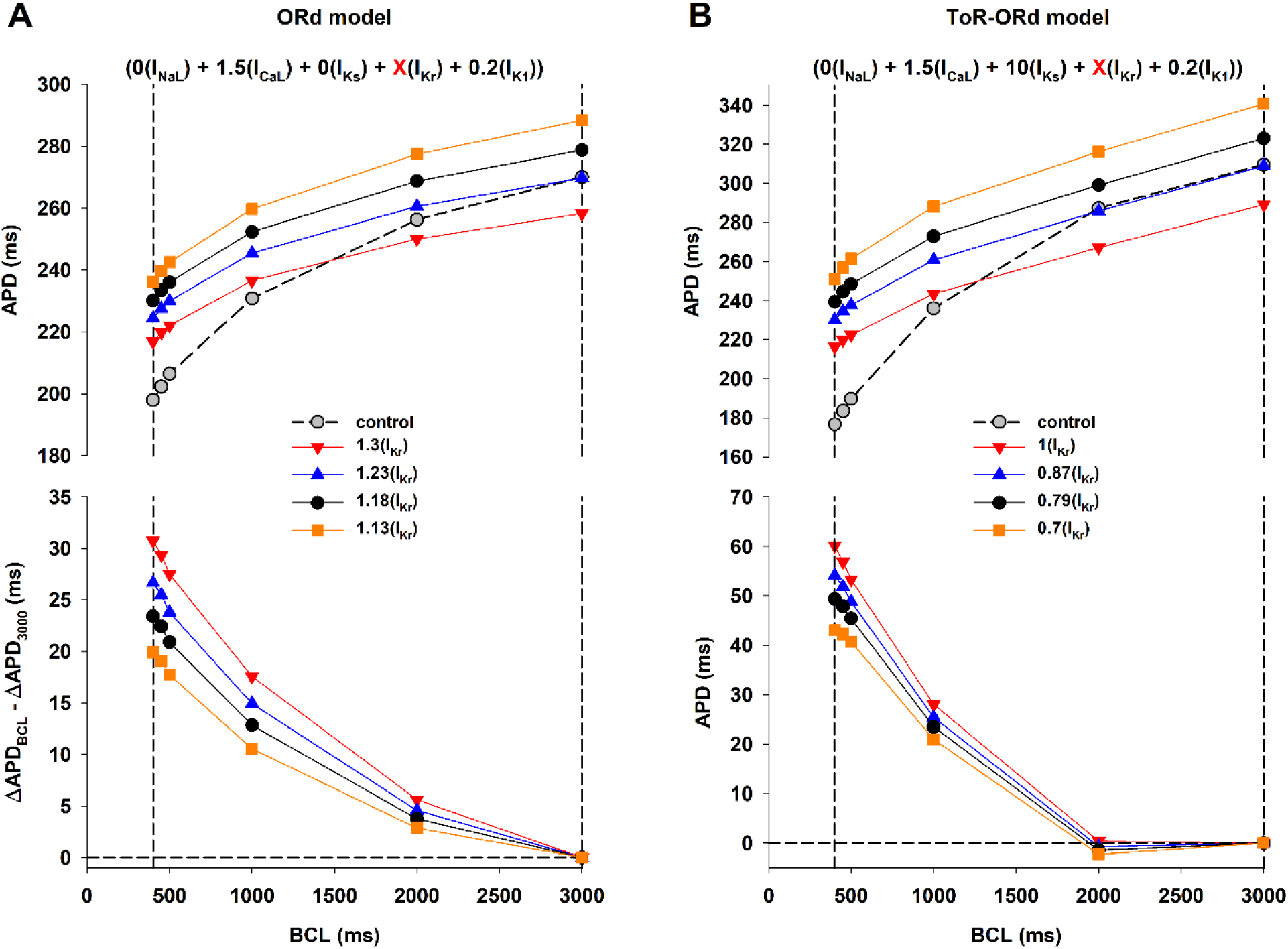
Interventions causing different amounts of APD prolongation with a positive rate dependence. Panel A (top): APD rate dependence with the ORd model for control (dashed black line) and four interventions with fixed modulation of I_NaL_, I_CaL_, I_Ks_ and I_K1_, and varying degrees of I_Kr_ modulation (0I_NaL_+1.5I_CaL_+0I_Ks_+XI_Kr_+0.2I_K1_), with X = 1.13 (ochre squares), 1.18 (black circles), 1.23 (blue triangles up) and 1.3 (red triangles down). Panel A (bottom): Difference between APD prolongation (with respect to control) for each BCL and APD prolongation with BCL = 3000 ms for the four interventions in Panel A (top). A positive value indicates positive rate dependence. The vertical dashed lines indicate BCL = 400 ms and 3000 ms. Panel B shows equivalent results with the ToR-ORd model using the same format. See text for detailed description.

The optimal combination of ion current modulation to achieve a positive rate dependent response requires the use I_Ks_ blockers and I_Kr_ activators in the ORd model (Figure 8A), but the use of I_Ks_ activators and I_Kr_ blockers in the ToR-ORd model (Figure 8B); the differences are a consequence of the different formulations of I_Ks_ and I_Kr_ in the models. Figure 9 shows a control action potential (Figure 9, top, dashed black line) and an action potential with a positive rate dependent response (Figure 9, top, solid blue line) having the same APD with BCL = 3000 ms (blue triangles up and gray circles in Figure 8). The figure shows that the dynamics of the combined effect of I_Ks_ + I_Kr_ during repolarization are similar for both models (Figure 9A and 9B, bottom): an increase of the delayed rectifier currents (i.e., I_Ks_ + I_Kr_) during phase 2 of the action potential and a decrease of the delayed rectifier currents during phase 3 are associated with a positive rate dependent response.

**Figure 9.**
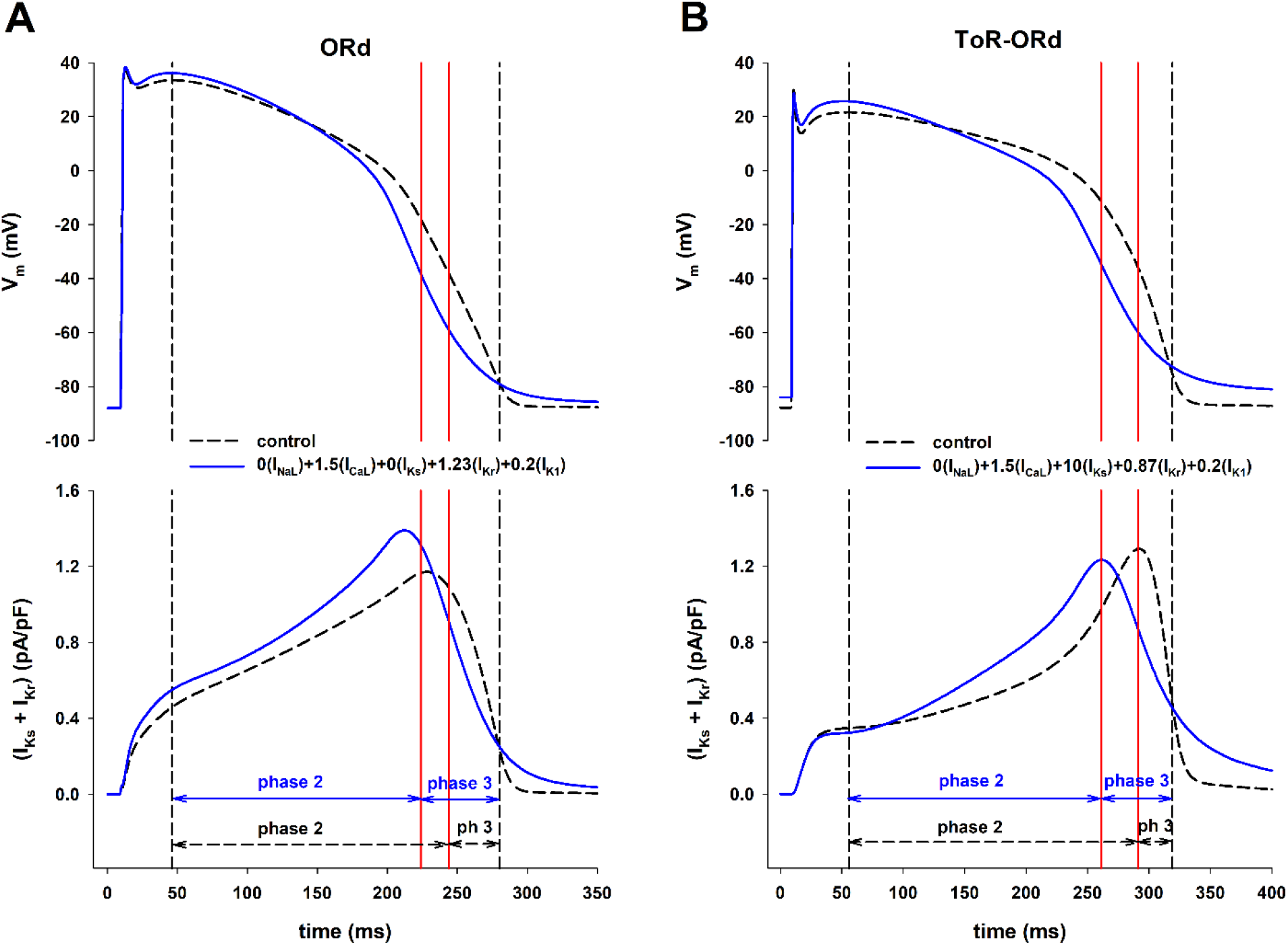
Dynamics of the total delayed rectifier potassium currents (= I_Ks_ + I_Kr_) for control action potentials and action potentials with positive rate dependence. Panel A (top): Control action potential (black dashed line) and action potential with positive rate dependence (solid blue line) with the same APD during stimulation with BCL = 3000 ms, with the ORd model. The leftmost vertical dashed line indicates the beginning of phase 2 repolarization; the rigthmost vertical dashed line indicates the time of 90% repolarization. The vertical red solid lines indicate the end of phase 2 (and beginning of phase 3) for both action potentials. Panel A (bottom): I_Ks_ + I_Kr_ during the action potentials in the top of panel A. The duration of phase 2 and 3 are indicated by the arrows. Panel B shows equivalent results for the ToR-ORd model, and it has the same format as panel A. See text for detailed description.

Figure 10 shows the repolarization reserve with BCL = 3000 ms for the interventions in Figure 8 with respect to the control action potential (gray bar, dashed horizontal line). For both models, the larger the APD prolongation (ochre squares in Figure 8A and 8B), the smaller the repolarization reserve (X=1.13I_Kr_, ochre bars in Figure 10A and 10B). Note that for interventions that prolong APD with BCL = 400 ms but shorten it with BCL = 3000 ms (red triangles down in Figure 8A and 8B), the repolarization reserve is just slightly decreased over that of control (X = 1.3I_Kr_ in Figure 10A and X = 1I_Kr_ in Figure 10B). The results in Figure 10 suggest that multichannel modulation of depolarizing and repolarizing currents that prolong APD at fast heart rates (close to those occurring during ventricular tachycardia) and shorten APD at slow heart rates should minimize the risk of triggered arrythmias.

**Figure 10.**
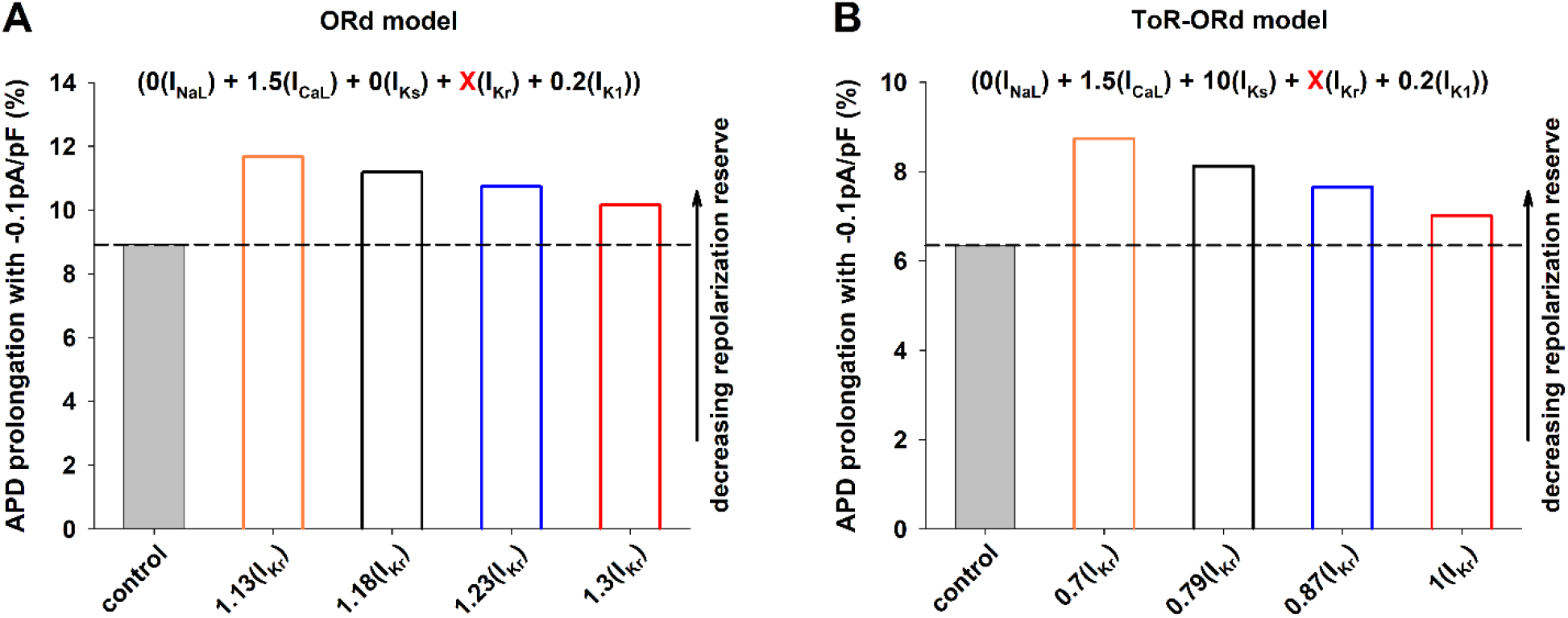
Percentage prolongation of APD when a constant depolarizing current of −0.1pA/pF is applied during the action potential when BCL = 3000 ms for the control action potential (gray bar, dashed line) and for the interventions in Figure 8. Panel A: Interventions that result in a positive rate dependence using the ORd model (0I_NaL_+1.5I_CaL_+0I_Ks_+XI_Kr_+0.2I_K1_, where X is the value in the x-axis). Panel B: Interventions that result in a positive rate dependence using the ToR-ORd model (0I_NaL_+1.5I_CaL_+10I_Ks_+XI_Kr_+0.2I_K1_, where X is the value in the x-axis). For all interventions, APD prolongation is above the dashed line indicating that the repolarization reserve is decreased with respect to control. See text for detailed description.

## DISCUSSION

We have shown that, in computer models of the human ventricular action potential, the combined modulation of both depolarizing and repolarizing ion currents (I_NaL_, I_CaL_, I_Ks_, I_Kr_ and I_K1_) results in a stronger positive rate dependent APD prolongation than modulation of just repolarizing potassium currents (I_Ks_, I_Kr_ and I_K1_); modulation of a single repolarizing current (I_Kr_) results in reverse rate dependence (Figures 1–3). A strong positive rate dependent APD prolongation correlates with an acceleration of phase 2 repolarization and a deceleration of phase 3 repolarization, which leads to a triangulation of the action potential shape (Figure 4). A positive rate dependent APD prolongation decreases the repolarization reserve at slow excitation rates (Figure 5), which can be managed by interventions that prolong APD at fast excitation rates and shorten APD at slow excitation rates (Figures 8 and 10). Feature importance analysis shows that, in both the ORd and ToR-ORd models, I_CaL_ and I_K1_ are the most important ion currents to achieve a positive rate dependent APD prolongation (Figures 6 and 7).

There is experimental and clinical evidence showing that selective block of a specific potassium current results in reverse rate dependent APD prolongation (Dorian and Newman 2000; Virag et al. 2009; Banyasz et al. 2009; Barandi et al 2010). There is also clinical and experimental evidence suggesting that the combined modulation of several ion currents may result in neutral (neither positive nor reverse) rate dependent APD prolongation (Hondeghem and Snyders 1990; Dorian and Newman 2000; Kassotis et al. 2003; Shibata et al. 2012). The exact mechanism underlying the attenuation of reverse rate dependence by the combined modulation of several ion currents is unknown. In that context, our results provide further understanding of how multichannel pharmacology can produce a robust positive rate dependent APD prolongation by optimal modulation of both depolarizing and repolarizing ion currents, using ion channel activators and blockers, which should result in an antiarrhythmic effect.

We have shown earlier (Cabo 2022) that the combination of potassium channel activators and blockers can produce a positive rate dependent APD prolongation in two different models of the human ventricular action potential: the TNNP model (ten Tusscher et al. 2004) and the ToR-ORd model (Tomek et al. 2019). Despite differences in the formulation of major repolarizing currents in the models (for example, the TNNP model has an unphysiologically large I_Ks_) both models predict a positive rate dependent APD prolongation by enhancing I_Ks_ and blocking I_Kr_ and I_K1_. In this report, we show that optimal modulation of those same three potassium currents also results in positive rate dependent APD prolongation in the ORd model (O’Hara et al. 2011). However, in contrast to the other two models, positive rate dependence in the ORd model requires complete block of I_Ks_ (instead of I_Ks_ enhacement), enhancement of I_Kr_ (instead of I_Kr_ inhibition) and I_K1_ block. Despite differences in the formulation of the ion currents, the dynamics of the total delayed rectifier current (i.e., I_Ks_ + I_Kr_) are similar in both models and further support the idea that an acceleration of phase 2 and deceleration of phase 3 repolarization of the action potential is associated with a positive rate dependent response (solid blue line in Figure 9).

In all models tested in this and earlier reports (Cabo 2022), I_K1_ block is important to achieve a positive rate dependent APD prolongation. Indeed, in two of the models (TNNP and ToR-ORd) (Cabo 2022), I_K1_ block alone can produce a positive rate dependent APD prolongation (see also Figure 7 for the ToR-ORd model), in agreement with an earlier simulation study (Cummins et al. 2014). In contrast, I_K1_ block does not result in positive rate dependence in the ORd model (Figure 6). Experimental evidence on whether I_K1_ inhibition results in positive or reverse rate dependence is mixed. Positive rate dependence has been demonstrated in guinea-pig myocytes treated with I_K1_ blocker terikalant (Williams et al. 1999), while inhibition of I_K1_ with either terikalant (Biliczki et al. 2005) or BaCl_2_ (Virag et al. 2009) in canine myocytes results in reverse rate dependence.

Chronic amiodarone and ranolazine (Belardinelli et al. 2006), which block I_NaL_, I_Ca_, I_Ks_ and I_Kr_, exhibit a neutral (neither positive nor reverse) rate dependent APD prolongation, in contrast with the reverse rate dependent APD prolongation of Class III agents that block a specific ion channel type like I_Kr_. This multichannel inhibition profile has been linked to their antiarrhythmic efficacy as well as their lack of proarrhythmic effects (Antzelevitch et al. 2004). The neutral rate dependent APD prolongation of amiodarone and ranolazine is associated with block of depolarizing and repolarizing ion currents, which has two opposing effects on APD: block of I_NaL_ and I_Ca_ tend to shorten APD; block of I_Ks_ and I_Kr_ tend to lenghten APD. In our simulations, a robust positive rate dependence response also requires modulation of depolarizing and repolarizing ion currents. The strong positive rate dependent APD prolongation shown in Figure 1 (black circles) is also associated with two opposing effects of depolarizing and repolarizing currents on APD, but, in contrast to amiodarone and ranolazine, those effects are achieved by the use of ion channel activators instead of ion channel blockers: enhancement of I_CaL_ tends to lengthen APD; enhancement of the total delayed rectifier current tends to shorten APD. The results in Figure 1 further suggest that modulation of multiple ion channels with activators and blockers (Figure 1, black circles) can produce a more robust positive rate dependence than modulation of those same ion channels with just ion channel blockers (Figure 1, green triangles down) at fast excitation rates (BCLs < 500 ms). I_Ks_ activators (Xu et al. 2002, Tamargo et al. 2004, Xu et al. 2015) and I_Kr_ activators (Shi et al 2020) have been developed to prevent excessive APD prolongation that may occur in patients suffering LQT syndrome, cardiac hypertrophy or cardiac failure. I_CaL_ activators have been shown to prevent reentrant arrhythmias in experimental models of myocardial infarction (Cabo et al 2000).

The multichannel inhibition profile of amiodarone and ranolazine has been linked to their antiarrhythmic efficacy as well as their lack of proarrhythmic effects (Antzelevitch et al. 2004). Block of depolarizing currents like I_NaL_ and I_CaL_ result in acceleration of phase 2 repolarization, preventing the initiation of triggered arrhythmias that may result in Torsade de Pointes. In our simulations, a positive rate dependent APD prolongation is also associated with an acceleration of phase 2 repolarization (Figure 4). However, the mechanism of that acceleration is different: the inhibition of I_NaL_ and the enhancement of the delayed rectifier currents (I_Ks_ + I_Kr_) compensate the enhancement of I_CaL_. While acceleration of phase 2 repolarization can reduce the likelihood of triggered arrhythmias, a strong positive rate dependent APD prolongation in our simulations is associated with a decrease in the repolarization reserve with respect to control action potentials (Figure 5), which could indicate a pro-arrhythmic risk. The decrease in the repolarization reserve can be managed by limiting the amount of I_K1_ inhibition. Interventions with moderate I_K1_ block (<= 50%) still result in a robust positive rate response with moderate decrease in the repolarization reserve with respect to control (Cabo 2022). Additionally, interventions that prolong APD at fast excitation rates, but shorten APD at slow excitation rates further moderate the decrease in the repolarization reserve and the risk of initiation of triggered arrhythmias (Figures 8 and 10).

In conclusion, multichannel modulation of depolarizing (I_NaL_ and I_CaL_) and repolarizing currents (I_Ks_, I_Kr_ and I_K1_), with ion channel activators and blockers, results in a robust APD prolongation at fast excitation rates, which should be anti-arrhythmic, while minimizing APD prolongation at slow heart rates, which should reduce potential pro-arrhythmic risks.

## Limitations

Computer models of the action potential integrate, often conflicting, experimental data obtained under different conditions from different preparations. As a consequence, conclusions derived from numerical simulations of the cardiac action potential should be interpreted with caution as predictions that need to be tested experimentally. The ORd and ToR-ORd models used in this report simulate healthy ventricular action potentials. But the density and kinetics of ion channels of myocytes is remodeled by disease (Baba et al. 2005), and the effect of drug agents on remodeled myocytes may be different from their effect in healthy myocytes (Cabo and Boyden 2003). Therefore, the conclusions of our simulations may not apply to diseased myocardial cells. Moreover, during the propagation of the action potential in myocardial tissue, electrotonic interaction between neighboring cells can modulate the effects of enhancing or inhibiting ion currents observed in single cells (Decker et al. 2009). Therefore, to elucidate how positive rate dependent APD prolongation affects the dynamics of propagation of premature impulses that initiate and sustain reentrant arrhythmias in heterogeneous and possibly remodeled myocardial tissue requires additional computational and experimental studies.

## REFERENCES

Antzelevitch C, Belardinelli L, Zygmunt AC, Burashnikov A, Di Diego JM, Fish JM, Cordeiro JM, Thomas G. Electrophysiological effects of ranolazine, a novel antianginal agent with antiarrhythmic properties. Circulation. 2004 Aug 24;110(8):904–10. doi: 10.1161/01.CIR.0000139333.83620.5D. Epub 2004 Aug 9. PMID: 15302796; PMCID: PMC1513623.

Baba S, Dun W, Cabo C, Boyden PA. Remodeling in cells from different regions of the reentrant circuit during ventricular tachycardia. Circulation 112, 2386–2396, 2005. doi:10.1161/CIRCULATIONAHA.105.534784

Banyasz T, Horvath B, Virag L, Barandi L, Szentandrassy N, Harmati G et al. Reverse rate dependency is an intrinsic property of canine cardiac preparations. Cardiovasc. Res. 84: 237–44, 2009.

Barandi L, Virag L, Jost N, Horvath Z, Koncz I, Papp R et al. Reverse rate-dependent changes are determined by baseline action potential duration in mammalian and human ventricular preparations. Basic Res Cardiol 105: 315–23, 2010.

Belardinelli L, Shryock JC, Fraser H. Inhibition of the late sodium current as a potential cardioprotective principle: effects of the late sodium current inhibitor ranolazine. Heart. 2006 Jul;92 Suppl 4(Suppl 4): iv6–iv14. doi: 10.1136/hrt.2005.078790. PMID: 16775092; PMCID: PMC1861317.

Biliczki P, Acsai K, Virag L, Talosi L, Jost N, Biliczki A, Papp JG, Varro A. Cellular electrophysiological effect of terikalant in the dog heart. European Journal of Pharmacology 510: 161–166, 2005.

Bloch-Thomsen P. DIAMOND (Danish Investigations of Arrhythmia and Mortality on Dofetilide). Clin Cardiol 21: 53–4, 1998.

Cabo C, Schmitt H, Wit AL. New mechanism of antiarrhythmic drug action: increasing L-type calcium current prevents reentrant ventricular tachycardia in the infarcted canine heart. Circulation. 2000 Nov 7;102(19):2417–25. doi: 10.1161/01.cir.102.19.2417. PMID: 11067798.

Cabo C, Boyden PA. Electrical remodeling of the epicardial border zone in the canine infarcted heart: a computational analysis. Am. J. Physiol. Heart Circ. Physiol. 284: H372–H384, 2003. doi:10.1152/ajpheart.00512.2002

Cabo C. Positive rate-dependent action potential prolongation by modulating potassium ion channels. Physiol Rep. 2022 Jun;10(12): e15356. doi: 10.14814/phy2.15356

Cummins MA, Dalal PJ, Bugana M, Severi S, Sobie EA (2014) Comprehensive Analyses of Ventricular Myocyte Models Identify Targets Exhibiting Favorable Rate Dependence. PLoS Comput Biol 10(3): e1003543. https://doi.org/10.1371/journal.pcbi.1003543

Decker, K. F., Heijman, J., Silva, J. R., Hund, T. J., & Rudy, Y. Properties and ionic mechanisms of action potential adaptation, restitution, and accommodation in canine epicardium. American journal of Physiology. Heart and Circulatory Physiology, 296(4): H1017–H1026, 2009. https://doi.org/10.1152/ajpheart.01216.2008

Dorian P, Newman D. Rate dependence of the effect of antiarrhythmic drugs delaying cardiac repolarization: an overview. Europace 2: 277–285, 2000.

Hondeghem LM, Snyders DJ. Class III antiarrhythmic agents have a lot of potential but a long way to go. Reduced effectiveness and dangers of reverse use dependence. Circulation 81: 686–90, 1990.

Hondeghem LM, Carlsson L, Duker G. Instability and triangulation of the action potential predict serious proarrhythmia, but action potential duration prolongation is antiarrhythmic. Circulation 103: 2004–2013, 2001.

Kannankeril P, Roden DM, Darbar D. Drug-Induced Long QT Syndrome. Pharmacol. Rev. 62: 760–781, 2010.

Kassotis J, Sauberman RB, Cabo C, Wit AL, Coromilas J. Beta Receptor Blockade Potentiates the Antiarrhythmic Actions of d-Sotalol on Reentrant Ventricular Tachycardia in a Canine Model of Myocardial Infarction. J. Cardiovasc. Electrophysiol. 14: 1233–1244, 2003.

Kennedy J, Eberhart R. “Particle Swarm Optimization”. Proceedings of IEEE International Conference on Neural Networks. IV. pp. 1942–1948, 1995. doi:10.1109/ICNN.1995.488968

Køber L, Thomsen PEB, Møller M, Torp-Pedersen C, Carlsen J, Sandøe E et al. Effect of dofetilide in patients with recent myocardial infarction and left-ventricular dysfunction: a randomised trial. Lancet 356: 2052–8, 2000.

O’Hara T, Virág L, Varró A, Rudy Y (2011) Simulation of the Undiseased Human Cardiac Ventricular Action Potential: Model Formulation and Experimental Validation. PLoS Comput Biol 7(5): e1002061. https://doi.org/10.1371/journal.pcbi.1002061.

Peters NS, Cabo C, Wit AL. Arrhythmogenic mechanisms: automaticity, triggered activity, and reentry. Cardiac electrophysiology: from cell to bedside. WB Saunders. 2000.

Shah RR, Hondeghem LM. Refining detection of drug-induced proarrhythmia: QT interval and TRIaD. Heart Rhythm 2: 758–772, 2005.

Shi YP, Pang Z, Venkateshappa R, Gunawan M, Kemp J, Truong E, Chang C, Lin E, Shafaattalab S, Faizi S, Rayani K, Tibbits GF, Claydon VE, Claydon TW. The hERG channel activator, RPR260243, enhances protective *I*_Kr_ current early in the refractory period reducing arrhythmogenicity in zebrafish hearts. Am J Physiol Heart Circ Physiol. 2020 Aug 1;319(2):H251–H261. doi: 10.1152/ajpheart.00038.2020. Epub 2020 Jun 19. PMID: 32559136.

Shibata S, Okamoto Y, Endo S, Ono K. Direct effects of esmolol and landiolol on cardiac function, coronary vasoactivity, and ventricular electrophysiology in guinea-pig hearts. J. Pharmacol. Sci. 118: 255–265, 2012.

Tamargo, J., Caballero, R., Gomez, R., Valenzuela, C., & Delpon, E. (2004). Pharmacology of potassium channels. Cardiovascular Research, 62, 9–33.

ten Tusscher K.H.W.J., Noble D., Noble P.J., and Panfilov A.V. A model for human ventricular tissue. Am. J. Physiol. Heart Circ. Physiol. 286: H1573–H1589, 2004.

Tomek J, Bueno-Orovio A, Passini E, Zhou X, Minchole A, Britton O, Bartolucci C, Severi S, Shrier A, Virag L, Varro A, Rodriguez B. Development, calibration, and validation of a novel human ventricular myocyte model in health, disease, and drug block. Elife 24;8: e48890, 2019. doi: 10.7554/eLife.48890.

Varro A, Baczko I. Cardiac ventricular repolarization reserve: a principle for understanding drug related proarrhythmic risk. Br. J. Pharmacol. 164: 14–36, 2011. doi: 10.1111/j.1476-5381.2011.01367

Virag L, Acsai K, Hala O, Zaza A, Bitay M, Bogats G, Papp JG, Varro A. Self augmentation of the lengthening of repolarization is related to the shape of the cardiac action potential: implication for reverse rate dependency. Br. J. Pharmacol. 156:1076–1084, 2009.

Waldo AL, Camm AJ, deRuyter H, Friedman PL, MacNeil DJ, Pauls JF et al. Effect of D-sotalol on mortality in patients with left ventricular dysfunction after recent and remote myocardial infarction. Lancet 348: 7–12, 1996.

Williams BA, Dickenson DR, Beatch GN. Kinetics of rate-dependent shortening of action potential duration in guinea-pig ventricle; effects of *I*_K1_ and *I*_Kr_ blockade. British Journal of Pharmacology 126: 1426–1436, 1999.

Xu X, Salata JJ, Wang J, Wu Y, Yan GX, Liu T, Marinchak RA, Kowey PR. Increasing *I*_Ks_ corrects abnormal repolarization in rabbit models of acquired LQT2 and ventricular hypertrophy. Am. J. Physiol. Heart Circ. Physiol. 283: H664–H670, 2002.

Xu Y, Wang Y, Zhang M, Jiang M, Rosenhouse-Dantsker A, Wassenaar T, Tseng GN. Probing binding sites and mechanisms of action of an I_Ks_ activator by computations and experiments. Biophys. J. 108: 62–75, 2015.

